# Protonation effects in protein-ligand complexes - a case study of endothiapepsin and pepstatin A with computational and experimental methods

**DOI:** 10.1101/2024.09.12.612797

**Authors:** Helge Vatheuer, Oscar Palomino-Hernández, Janis Müller, Phillip Galonska, Serghei Glinca, Paul Czodrowski

## Abstract

Protonation states serve as an essential molecular recognition motif for biological processes. Their correct consideration is key to successful drug design campaigns, since chemoinformatic tools usually deal with default protonation states of ligands and proteins and miss atypical protonation states. The protonation pattern for the Endothiapepsin/PepstatinA (EP/pepA) complex is investigated using different dry lab and wet lab techniques. ITC experiments revealed an uptake of more than one mole of protons upon pepA binding to EP. Since these experiments were performed at physiological conditions (and not at pH=4 at which a large variety of crystal structures is available), a novel crystal structure at pH=7.6 was determined. This crystal structure showed that only modest structural changes occur upon increasing the pH value. This lead to computational studies to reveal the exact location of the protonation event. Both computational studies could reveal a significant pKa shift resulting in non-default protonation state and that the catalytic dyad is responsible for the uptake of protons. This study shows that assessing protonation states for two separate systems (protein and ligand) might result in the incorrect assignment of protonation states and hence incorrect calculation of binding energy.

## 2 Introduction

The correct determination of protonation states in a protein-ligand complex is crucial, since ionizable amino acids play a key role in the pH-dependent structure, binding interactions and catalysis.[1, 2, 3] From a protein perspective, 7 of the 20 proteinogenic amino acids contain an ionizable group, which is about 1/3 of the amino acid pool found in natural proteins. The charge state of these amino acids can be modulated either by the protein environment or by a bound ligand, or the protein environment can strongly influence ligands with ionizable groups. This in turn can influence the strength of drug binding or the structural properties of the protein.

From a ligand perspective, rational design of charge-assisted contacts between a protein and its ligand, guided by p*K*_a_ calculations of the respective complex has been shown to improve affinity, e.g. for a series of *lin*-benzoguanines binding to tRNA-guanine transglycosylase.[4] Moreover, the uncharged form of a drug can cross biological membranes more easily than its corresponding charged form. One approach could be to design a molecule that remains uncharged during transport across membranes, but becomes charged by proton release upon binding to the target through electrostatic interactions.[4]

One prominent example of modulated protein charge states is HIV-1 protease (PR). HIV-1 PR is an aspartic protease (AP) and its catalytic dyad is made up of two aspartic acids. It has its pH optimum range at 4.0–6.0, at which one of these aspartates is protonated whereas the other aspartate is deprotonated.[5] If the pH value is lowered to more acidic conditions, both aspartates become protonated. Thus, different pH values can alter the protonation states of ionizable residues, which can lead to a conformational change on either the protein or the protein-ligand complex.[6]

To reveal the protonation state in a protein-ligand complex we can employ methods such as high resolution X-ray crystallography, or a combination of high resolution X-ray and neutron diffraction data (Figure 1, or isothermal titration calorimetry (ITC). The first two methods are capable of resolving structures with an estimated standard deviation of 0.02 Å in bond length, allowing the distinction between a C-O single bond (1.34 Å) of a carboxyl group with a proton attached and a C=O double bond (1.20 Å) of the carboxyl side chain of an aspartate.[7, 8] Moreover, the addition of neutron diffraction data improves the data-to-parameter ratio, and allows for the hydrogens to also be resolved.[9] The second method, ITC, is regarded as the gold standard for the investigation of intermolecular interactions in solution.[10] In addition to obtaining thermodynamic parameters such as enthalpy and equilibrium constant (from which changes in free energy and entropy can be calculated) this method is able to determine the overall changes in the protonation setup during the binding reaction.[11] This can be achieved by performing measurements in different buffers with different ionization enthalpies, and these protonation events have been demonstrated for a variety of enzyme classes, including aspartic proteases, serine proteases and methyltransferases.[12, 4, 13] However, ITC is not capable to retrieve the exact (at an atomistic level) localization of the protonation events.

**Figure 1:**
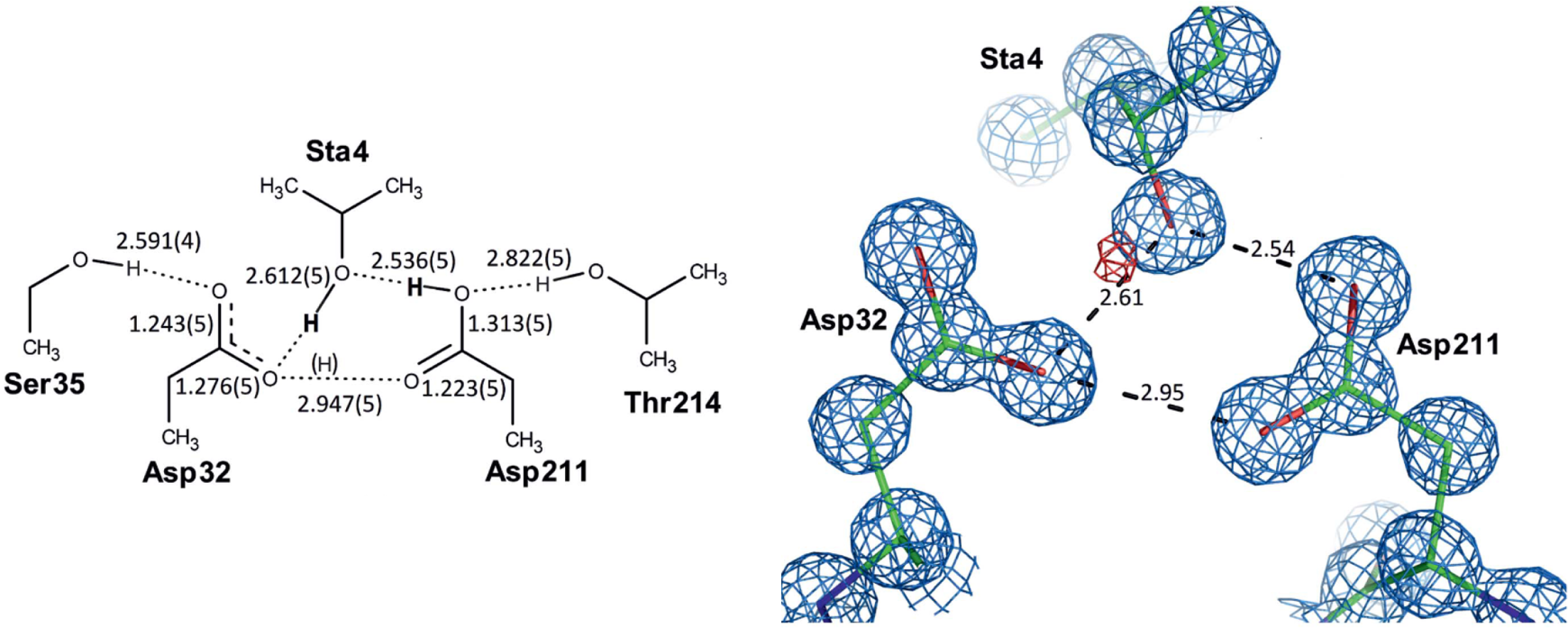
Polar interactions in the active site of the aspartic protease Sapp2p structure. *Left* : Schematic diagram with bond distances (values are in Å; estimated standard deviations of the distances are in parentheses). H atoms in bold are unambiguously assigned (the hydroxyl H atom of pepA was visible in the difference electron density; the H atom on the protonated carboxyl of Asp211 was deciphered from the C-O bond distances). The H atom in brackets is hypothetical. *Right* : Detailed structure of the active site in rod representation (PDB code: 4Y9W); hydrogen bonds are shown as dotted lines (numbers represent distances in Å). The 2F_o_-F_c_ electron density map contoured at the 1.5 level is shown in light blue; the F_o_-F_c_ difference electron density map contoured at the 2 level is shown in red. Figure modified from [8]

In this work, we use Endothiapepsin (EP), an exemplary aspartic protease, as a model enzyme to study protonation effects associated with ligand binding. The present protonation effect in the interaction of EP with the pan-protease inhibitor pepA was demonstrated by thermodynamic investigations using ITC measurements. Our results are compared with a published ITC study [12] and lead to new experiments such as crystal structure determination at physiological pH and novel computational approaches. These new experiments underline our experimentally (via ITC) determined protonation event upon pepA/EP complexation.

## 3 Results and Discussion

### 3.1 ITC measurements reveal proton transfer events

The total proton exchange (Δ*n*_H+_) in a binding event can be determined by ITC measurements in different buffers with distinct heats of ionization (ΔH_ionization_), according to the formula 1:

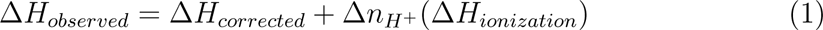

Integrating the raw thermograms yields the experimentally observed enthalpies (ΔH_observed_), which can be used for a linear regression with the slope corresponding to Δ*n*_H+_ and the y-axis intercept corresponding to the buffer-corrected enthalpy of the binding reaction (ΔH_corrected_).

The ITC measurements investigating the interaction of pepA and EP were car-ried out in in phosphate, ADA, HEPES and TRIS buffers at pH 7.0 with ranging ΔH_ionization_ from 3.60 to 47.45 kJ mol^−1^(Figure 2). From the overlay of the raw thermograms we observe a strong buffer dependency. It is clear that the Δ*n*_H+_ is not equal to zero, otherwise the raw thermograms would be congruent with each other (ΔH_obs_ = ΔH_corrected_).

**Figure 2:**
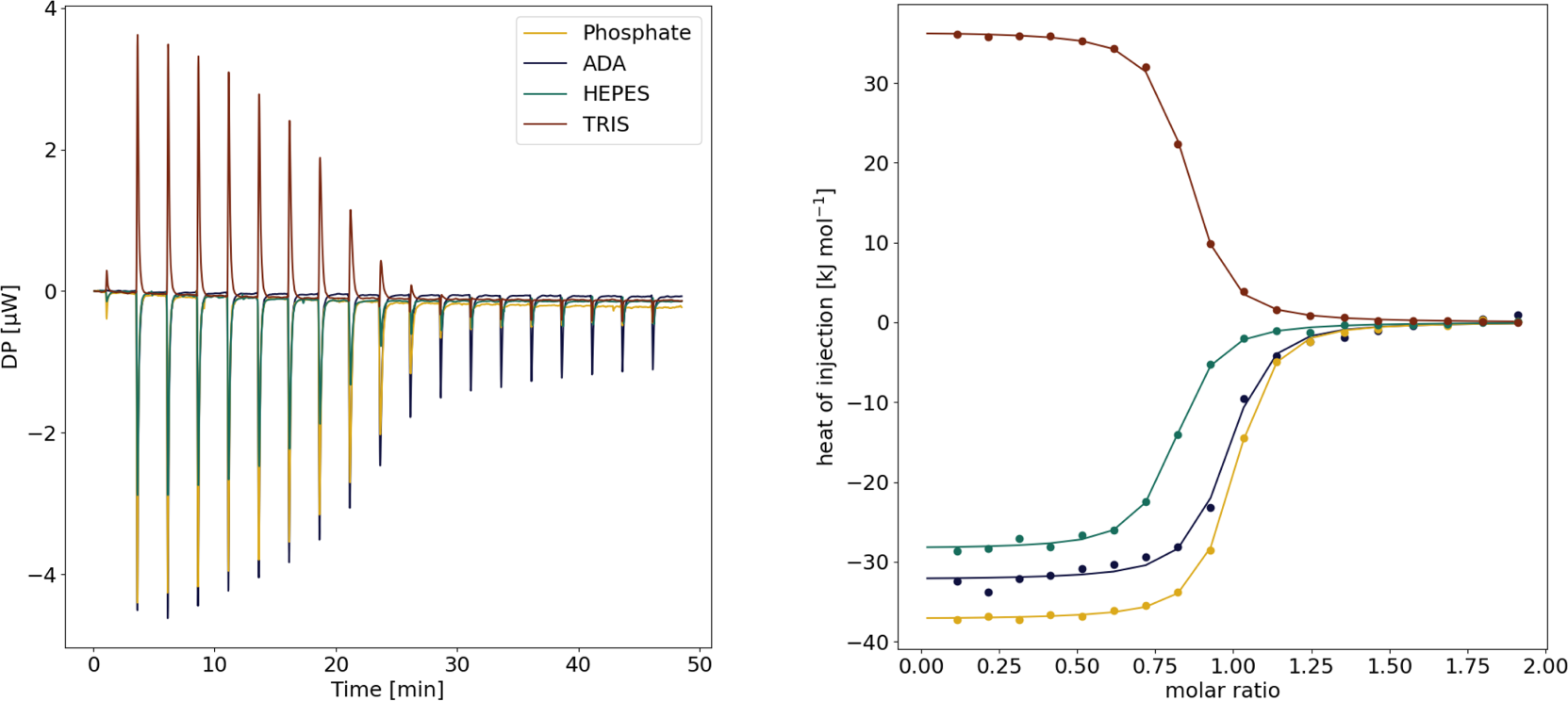
Overlay of raw thermograms of the binding reaction between EP and pepstatin A measured in phosphate, ADA, HEPES and TRIS at pH 7.0 (left); integrated data of the heat signals observed for the measurements in the four different buffers (right).

The ITC measurements indicated an uptake of 1.67 ± 0.12 protons per binding reaction (Figure 3). The TRIS outlier can be explained by its high positive ionization energy (47.45 kJ mol^−1^), which leads to a positive enthalpic contribution (30.7 ± 2.9 kJ mol^−1^) due to the proton uptake of the buffer.

**Figure 3:**
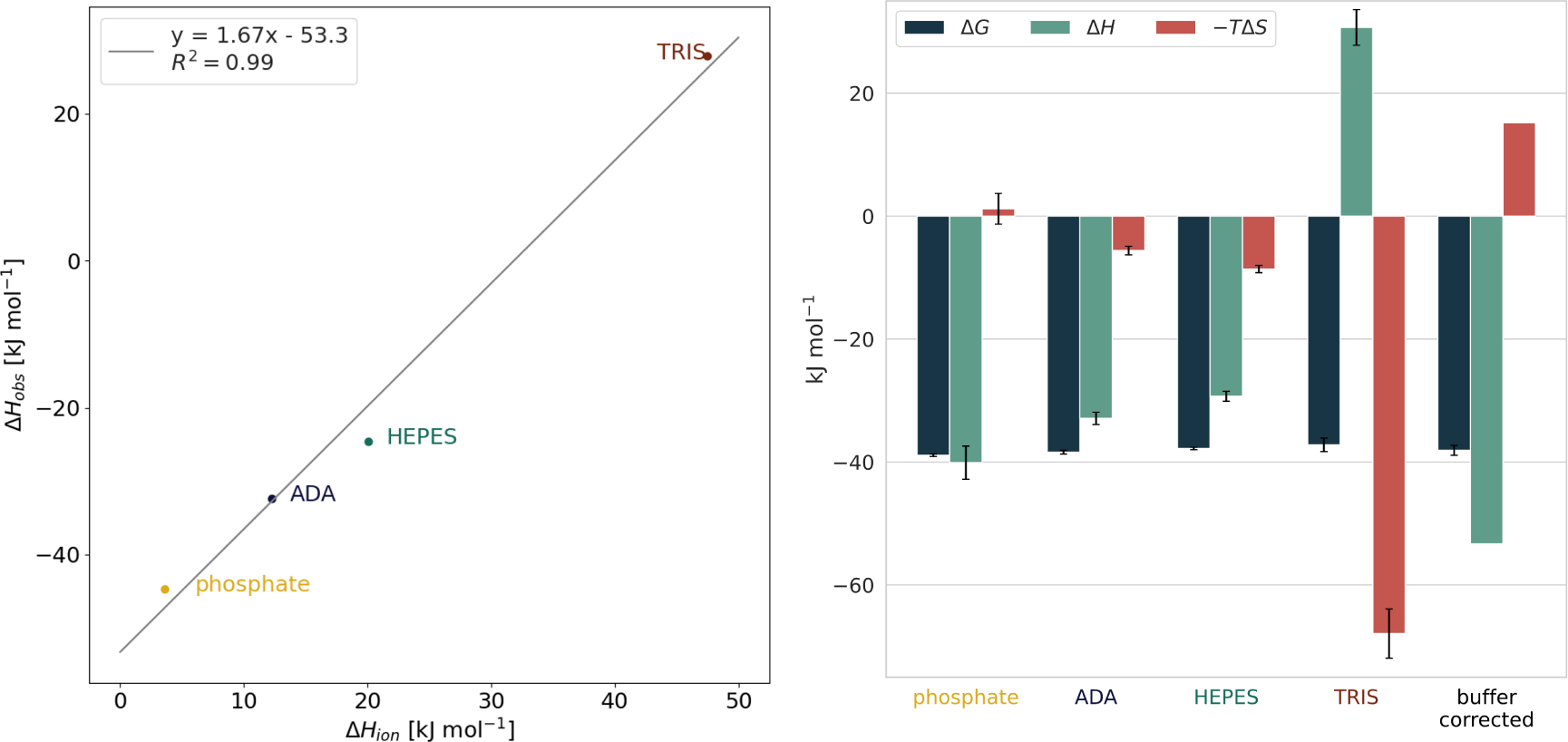
Left: Calculation of the heat of ionization. The experimentally observed enthalpies ΔH_obs_ are plotted against the heat of ionization ΔH_ion_ of the respective buffers (phosphate: 3.60 kJ mol^−1^ ADA: 12.23 kJ mol^−1^, HEPES: 20.04 kJ mol^−1^, TRIS: 47.45 kJ mol^−1^[14]). The slope of the line describes the proton uptake during the formation of the protein-ligand complex (on average 1.67 ± 0.12 mol), while its intersection with the ordinate describes the buffer-corrected enthalpy of the binding reaction (ΔH_corrected_ = – 53.3 kJ mol^−1^). Right: Thermodynamic profiles of complex formation in phosphate, ADA, HEPES and TRIS buffers and the buffer-corrected thermodynamic profile. For the buffer-corrected profile, the change in Gibbs free energy ΔG is calculated as the average of ΔG observed in the four buffers, ΔH is obtained as described above, and the entropic contri-bution –TΔS is calculated from the numerical difference between ΔG and ΔH.

This deviates - at least in terms of the absolute value - with the Δ*n*_H+_ previously published values of this interaction: 1.1 ± 0.1 protons revealed by ITC by Gómez and Freire.[12] In the following, we will compare our ITC setup with that one of Gómez and Freire, the major differences are given in Table 1.

**Table 1:** Experimental conditions used here compared to those used by Gómez and Freire[12].

| Parameter | This study | Literature |
| --- | --- | --- |
| c[EP] | 50 $\mu\text{M}$ | 80 $\mu\text{M}$ |
| c[pepA] | 500 $\mu\text{M}$ | 1200 $\mu\text{M}$ |
| c[Buffer] | 100 mM | 20 mM |
| Temperature | 25°C | 16°C |
| pH | 7.0 | 7.0 |
| DMSO (v/v) | 2.5 % | - |
| Tween20 (v/v) | 0.1 % | - |
| Cell volume | 0.2 mL | 1.36 mL |
| Used buffers | ADA, HEPES,<br>Phosphate, TRIS | ACES, Cacodylate, HEPES,<br>Phosphate, TRIS |

Cacodylate was excluded as a buffering agent due to its toxicity. Measurements in ADA buffer were only performed here, while we did not perform measurements in ACES buffer. PepA was applied from a 20 mM DMSO stock solution, resulting in a final concentration of 2.5 % (v/v). This changed parameter could be a reason for the different results, since it has been shown for several protein-ligand complexes that a higher concentration of DMSO in the samples leads to a decrease in affinity, even at a low concentration of 1 % (v/v).[15] 2.5 % (v/v) DMSO corresponds to a molar concentration of 0.35 M.

Furthermore, different experimental conditions could have affected the proton trans-fer:

- higher temperature
- concentration of the reagents and buffer/salt
- the presence of DMSO as solvent (already described in the previous paragraph)
- the calorimetric device used

Each parameter alone can have a clear effect on the thermodynamic properties.[11] We used the MicroCal PEAQ-ITC Automated ITC with a cell volume of 200 µL 25°C is a standard temperature, where most of the experiments are carried out, not only ITC. We followed standard protocols for protein, ligand, and buffer concentrations.[16]

Another parameter influencing the thermodynamic properties with respect to *K*_d_, Δ*G*, Δ*H* and *−T* Δ*S* is the protein concentration used. For a ligand binding to thermolysin with an affinity in the low micromolar range, the determined *K*_d_ in-creases with protein concentration by an order of magnitude from 0.86 µM at the lowest protein concentration of 50 µM to over 14 µM at 300 µM thermolysin.[11]

The affinity of pepA is pH dependent and becomes higher at more acidic pH, with a *K*_d_ < 10 nM for pH ≤ 5, reaching the upper detection limit of the ITC instrument. This could be another explanation for the deviation between our ITC results and those published by Gómez and Freire[12].

The stoichiometry of the EP-pepA binding reaction is illustrated in Figure 4. There is a noticeable difference between the stoichiometry in the respective buffers. Assuming that pepA was taken from the same stock solution, *i.e.* the ligand concentration is correct, this could indicate that EP is more unstable/less active in HEPES: over 90 % activity (*N ≥* 0.9) in phosphate versus 70% activity (*N ≈* 0.9) in HEPES, since crystallization and previous calorimetric measurements show a 1:1 binding of the ligand to the active site of the protein. At least in TRIS, the results suggest an increasing degradation of the protein: *N* decreased from above 0.8 to below 0.6 after about 10 hours of exposure at room temperature (RT), according to the batchwise measurements in the automated ITC device. It should be noted that the ITC is able to cool the samples in the tray prior to titration, but there have been no stability problems with EP at RT. We assume that the different stoichiometries should not affect the overall protonation event: it will be constant throughout different measured stoichiometries.

**Figure 4:**
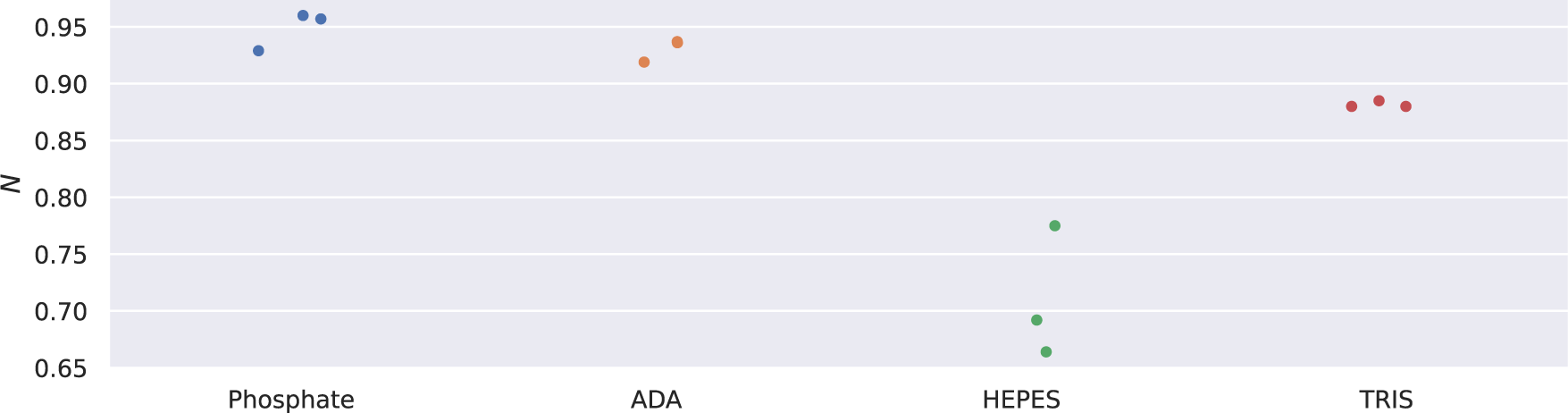
Overview of stoichiometry *N* of ITC measurements of the binding reaction between EP and pepA in different buffers of 100 mM concentration at a pH of 7.0.

However, EP as fungal enzyme has its optimum at an acidic pH, and most of the published experimental data, whether crystallographic or calorimetric, have been collected almost exclusively in an acetate buffer at pH 4.5–5.0, except for the study by Gómez and Freire.[12] Thus, pH together with temperature may have caused the decrease in activity of EP.

The available EP crystal structures in the PDB are exclusively available at an acidic pH of 4.5–5.0 due to the stability of EP. In contrast, the ITC experiments were performed at pH 7.0 which might have affected protein conformational changes. Therefore, we aimed for the determination of EP X-ray at pH values in the neutral range. Additionally, we used two different computational approaches to interprete the ITC data from a structural point of view, because the ITC data itself does not reveal the exact atomistic location of the protonation event.

### 3.2 Novel crystal structures at higher pH values reveal no major conformational changes

In order to have a structure at the same pH as the calorimetric measurements, EP and pepA were co-crystallized and soaked for 24 hours in soaking conditions with buffers of pH 7.6 and analyzed. The resolution was 1.23 Å, higher than the published 2.0 Å structure (PDB entry: 4ER2), and the all-atom RMSD between the pH 7.6 structure and 4ER2 amounts to 0.89 Å(Figure 5). However, the biggest variation within the binding site was located at both termini, where the α-carbon of the the isovaleryl residue at the N-terminus and the C-terminal α-carbon of statine is “flipped” (pepA sequence: Iva-Val-Val-Sta-Ala-Sta). This is because soaking at pH 7.6 results in a modified orientation compared to the structure in the acidic range.

**Figure 5:**
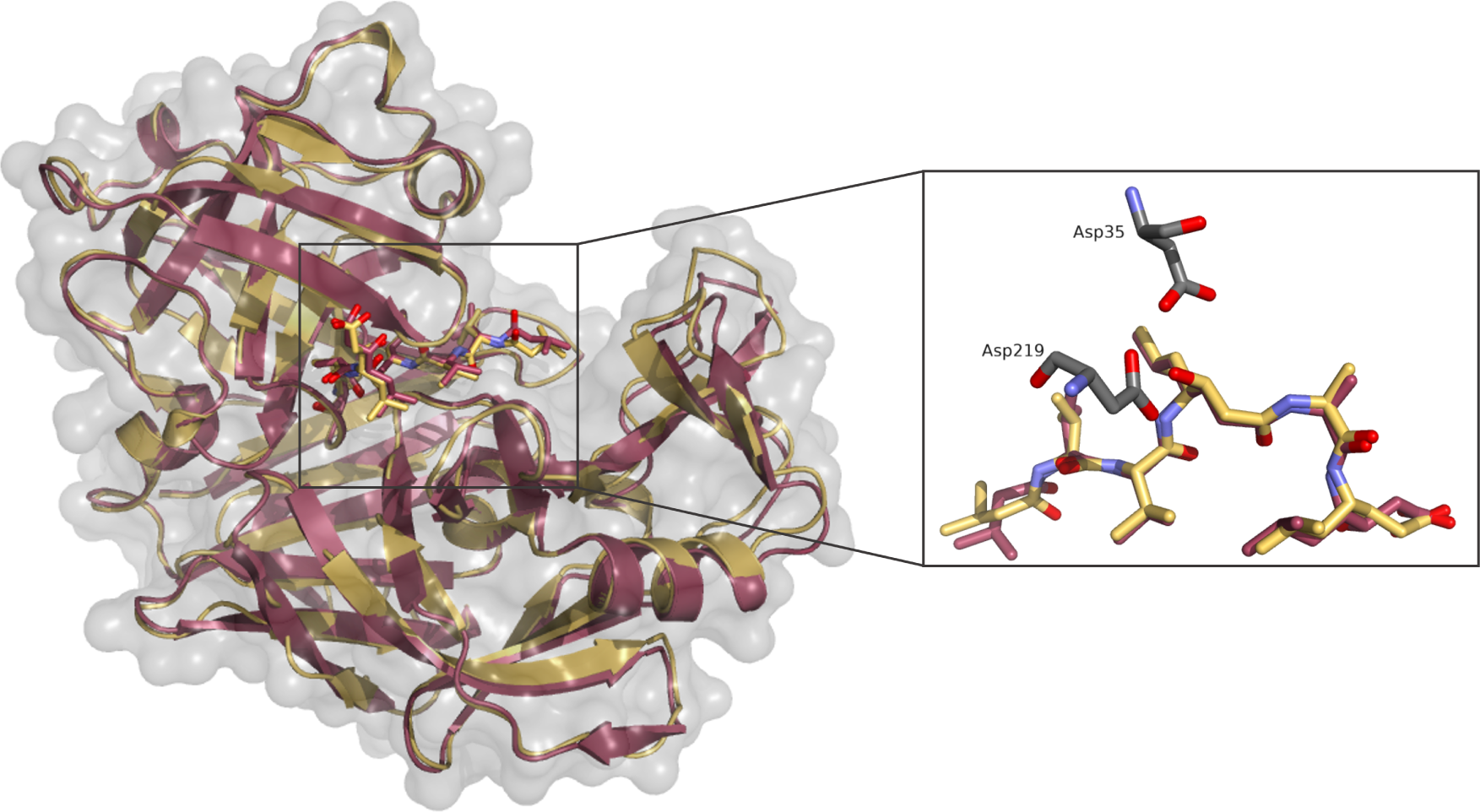
Comparison of EP-pepA complexes crystallized at pH 4.6 (red, PDB entry 4ER2) and soaked at pH 7.6 (yellow) with detailed view of pepA in the binding site.

Although there is a tiny residual density for the “acidic” conformation at pH 7.6 (i.e. a second conformation is observed for the crystal structure at different pH values) and vice versa, these are negligible.

### 3.3 Implicit solvent p*K*_a_ calculations allow for a structural interpretation

As a starting point, we used HIV-1 protease (HIV-1 PR) to calibrate our Poisson-Boltzmann (PB) methodology. The p*K*_a_ values resulting from the protein p*K*_a_ calculations for HIV-1 PR demonstrate that our calculations are in very good agreement to the experimental values: while Asp25 carboxylic side chain is the more acidic one with a p*K*_a_ of 3.2, Asp25’ carboxylic group has a p*K*_a_ of 5.7. The corresponding experimental values are 3.1 and 5.2 (Table 2).

**Table 2:** Results of the p*K*_a_ calculations on HIV-1 PR (PDB entry: 1XL5)

| **Method** | **Asp25** | **Asp25'** |
| --- | --- | --- |
| Experiment[17] | 3.1 | 5.2 |
| ZAP-based PBS | 3.2 | 5.7 |

The structural similarity between HIV-1 PR and EP in the active site is evident, the distances between the aspartates are almost identical (Figure 6). This convinced us to apply the HIV-1-PR validation protocol also for EP.

**Figure 6:**
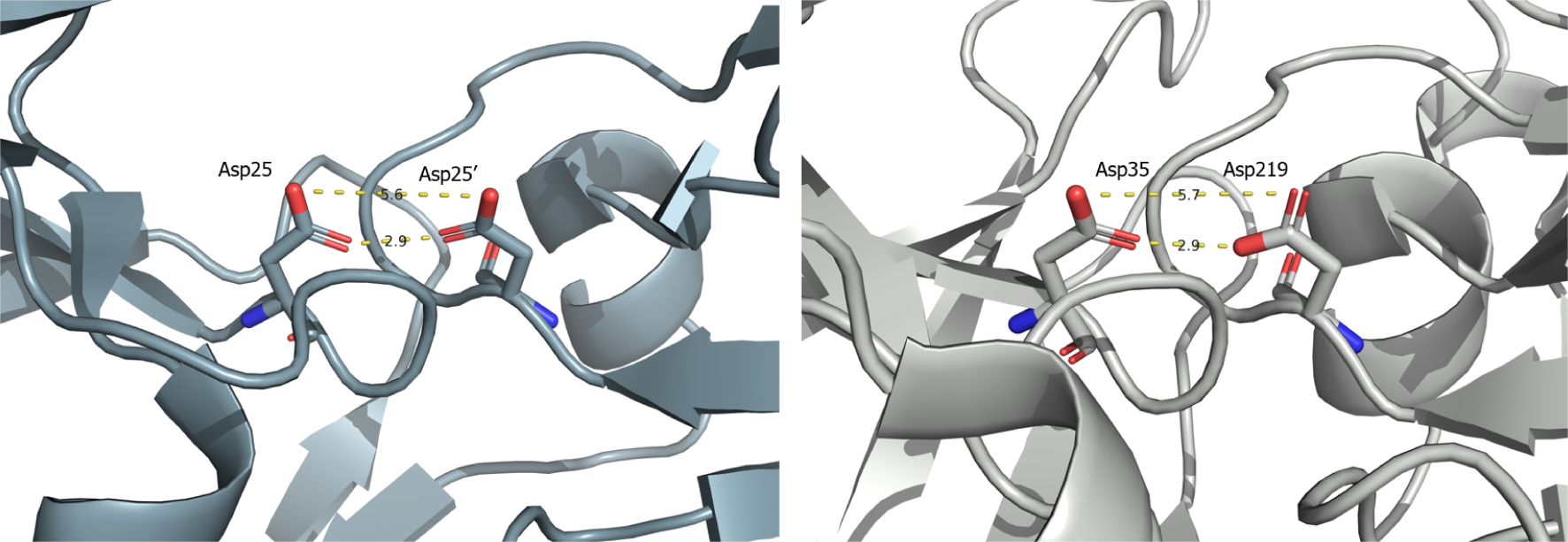
Comparison of the active site of HIV-1 PR ligand-deleted (left, PDB entry: 4EJ8) and EP (right, 4Y5L) apo structures. Distance between the relevant atoms in Å.

Next, we carried out p*K*_a_ calculations with our PB solver on the three different EP-pepA structures (Table 3). The C-terminal carboxy group of pepA was titrated analogous to the titratable amino acid side chains. The only slight p*K*_a_ shift of the pepA carboxy group can be explained by the solvent exposure and the associated lack of interactions with amino acid side chains. The overall exchange of protons transferred during the binding reaction (Δ*n*_H+_) is 0.79 for the structure 4ER2 and 0.5 for the pH 7.6 structure. In all three structures, the three amino acids Asp33, Asp35 and Asp 219 are mainly responsible for the shift. While Asp81 shows a delta Δp*K*_a_ of 0.5 to 0.6, it does not lead to a protonation shift at an assumed pH of 7.0, since the aspartate is more acidic than the other three. The calculated Δ*n*_H+_ of the complex at pH 7.6 was 0.5, probably due to the structural changes mentioned above.

**Table 3:** Results of the p*K*_a_ calculations on the EP-pepA complexes crystallized and afterwards soaked at different pH and the resulting protonation change Δ*n*_H+_ (in mols) at pH 7.0 (the pH value of the ITC experiments). The ligand p*K*_a_ is the C-terminal carboxy group.

| Crystal pH | | Asp35 | Asp219 | Asp33 | Asp81 | Ligand | $\sum \Delta n_{H^+}$ (pH 7.0) |
| --- | --- | --- | --- | --- | --- | --- | --- |
| 4.6 | Ligand-deleted | 5.8 | 5.5 | 6.3 | 4.6 | 4.32 | 0.79 |
|  | Complex | 6.4 | 7.0 | 6.8 | 5.2 | 4.4 |  |
| | $\Delta n_{H^+}$ | 0.14 | 0.47 | 0.22 | 0.02 | 0 | |
| 7.6 | Ligand-deleted | 6.2 | 5.5 | 6.4 | 4.6 | 4.32 | 0.5 |
|  | Complex | 6.3 | 7.0 | 6.7 | 5.1 | 4.2 |  |
| | $\Delta n_{H^+}$ | 0.03 | 0.47 | 0.13 | 0.01 | 0 | |

A closer look was then taken at the binding pocket with pepA in the active site of EP, and in particular at the interacting aspartate residues (Figure 7). The structures differ by a maximum of 0.2 Å. The most prominent interacting groups are the carboxylic side chain of both Asp35 and Asp219, the catalytic dyad, with the hydroxyl group of statine 4 of pepA. While the distance between the side chains oxygens of Asp33 and Asp35 is too high to form a hydrogen bond (6.2 Å), one oxygen is in distance of 3.6 Å to the acylic side chain of the pepA ligand. This interaction is probably responsible for the slight p*K*_a_ shift of Asp33.

**Figure 7:**
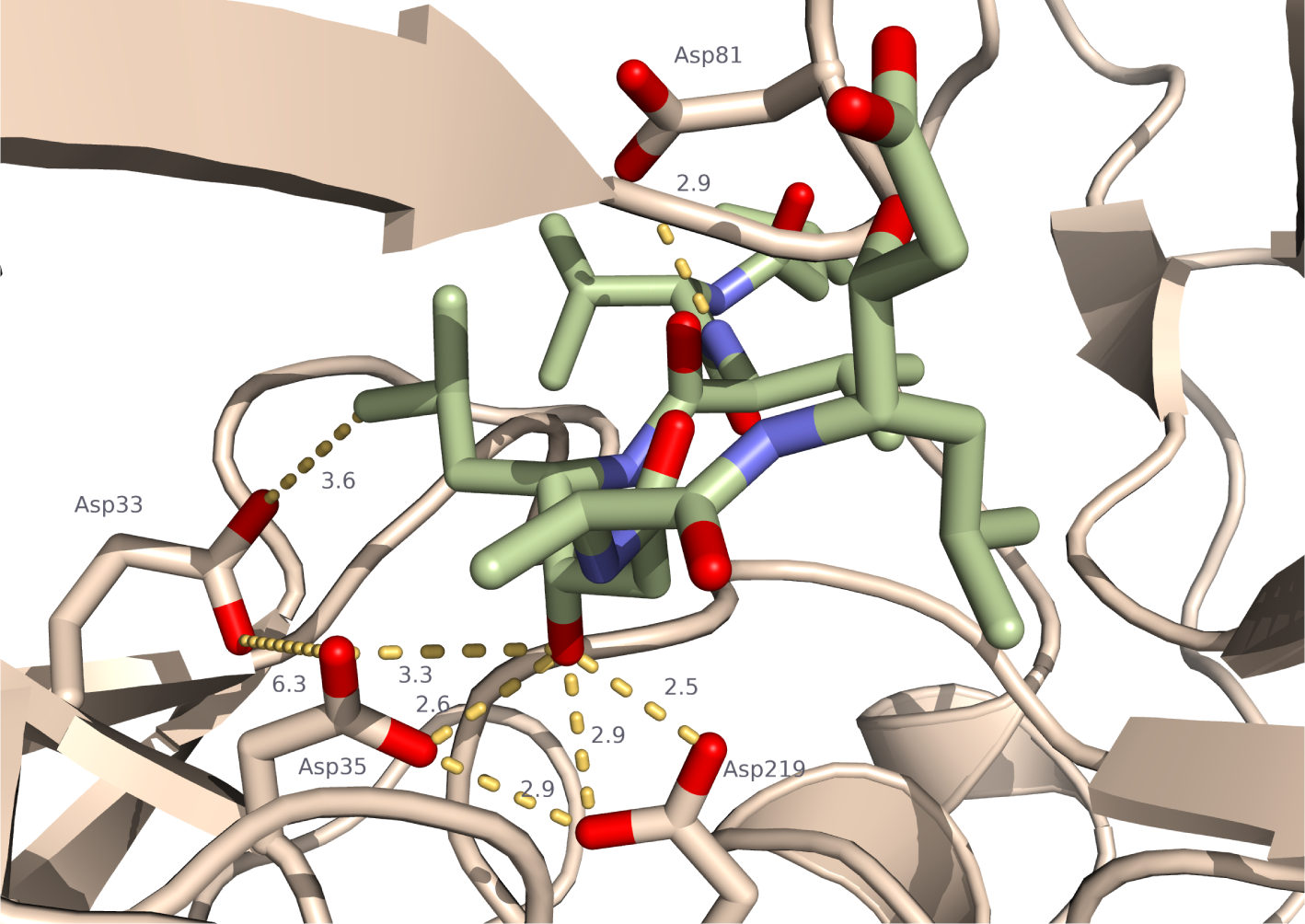
pepA (light green) in the active site of EP. The three carboxylic side chains of Asp33, Asp35 and Asp219 are the main contributors to the protonation effect revealed with PB solver. Asp81 is the fourth Asp within the binding site, but not part of the protonation effect according to the calculations. Distance between the relevant atoms in Å (PDB entry: 9GFY - pH 7.6 structure).

Most of the protonation effect based on our calculations occurs at the catalytic dyad. Since Asp35 and Asp219 behave like a dicarboxylic acid due to the given structural conditions and the small distance between them, Asp219 is most likely protonated at pH 7.0, whereas no change for the catalytic dyad occurs at pH 4.6 during complex formation (Figure 8).

**Figure 8:**
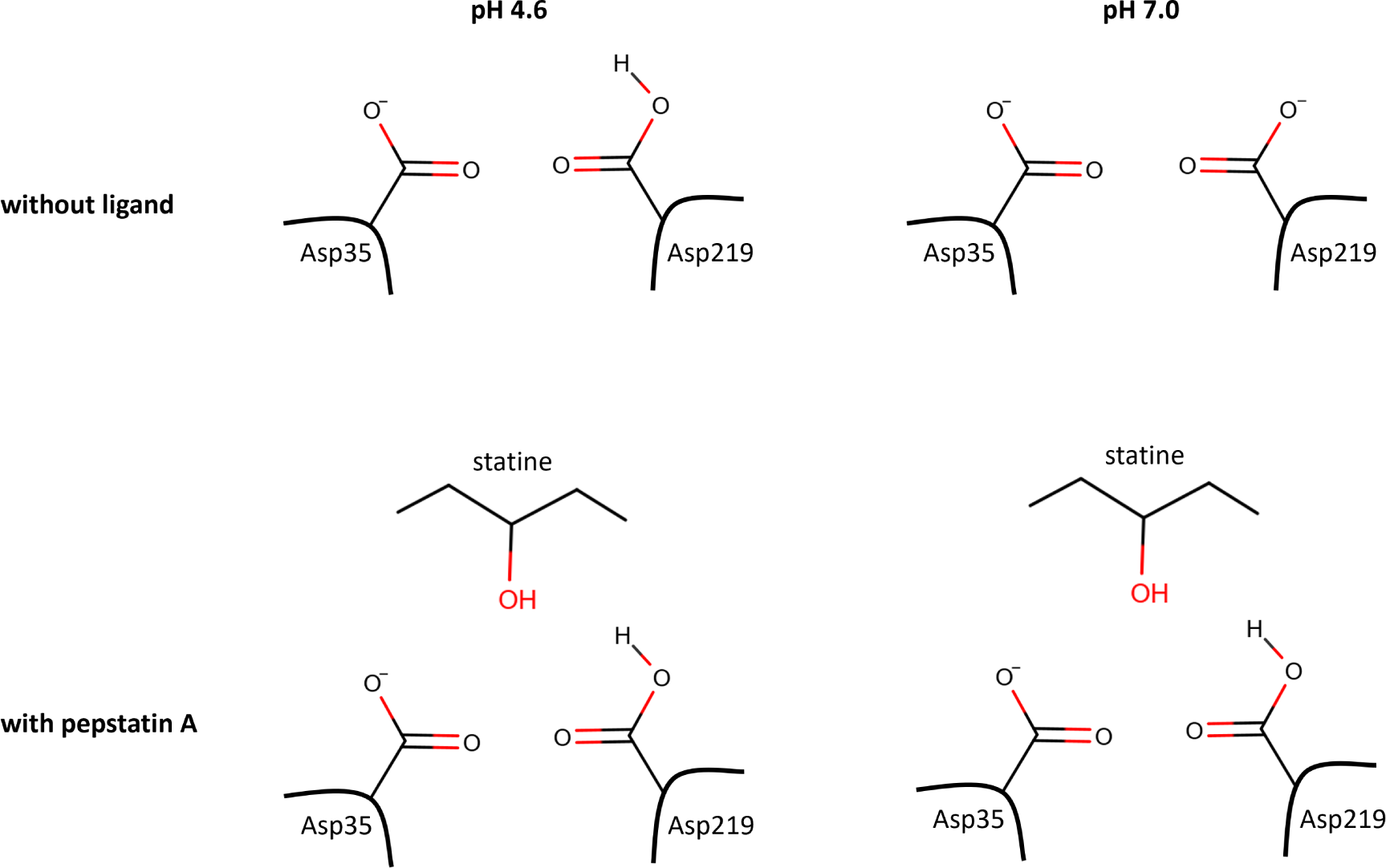
Overview of the hypothesized protonation states of the catalytic dyad of EP without ligand at pH 4.6 and 7.0, and with pepA at both pH values. At pH 4.6 the catalytic dyad is monoprotonated without ligand and with pepA, while at pH 7.0 both Asp are deprotonated, but Asp219 takes a proton upon binding of pepA.

### 3.4 Constant-pH MD simulations further underline the observed proton transfer event

Intrigued by the above results, we set out to understand the behavior of the selected aspartates through constant pH molecular dynamics (CpHMD) simulations. CpHMD is well-suited for investigating molecular systems characterized by pH-dependent dynamics and functions.[18] Thus, we performed simulations in both apo and pepA-bound EP at different pH values.

From a structural point of view, the root mean square deviation (RMSD) of apo and pepstatin-bound EP with respect to the crystal structure of 4ER2 do not show a strong dependence on the pH of the environment (Figure 9). Overall, unliganded EP shows higher RMSD values that fluctuate strongly. In comparison, pepstatin-bound EP shows remarkably lower RMSD values, as well as a reduced amount of fluctuations.

**Figure 9:**
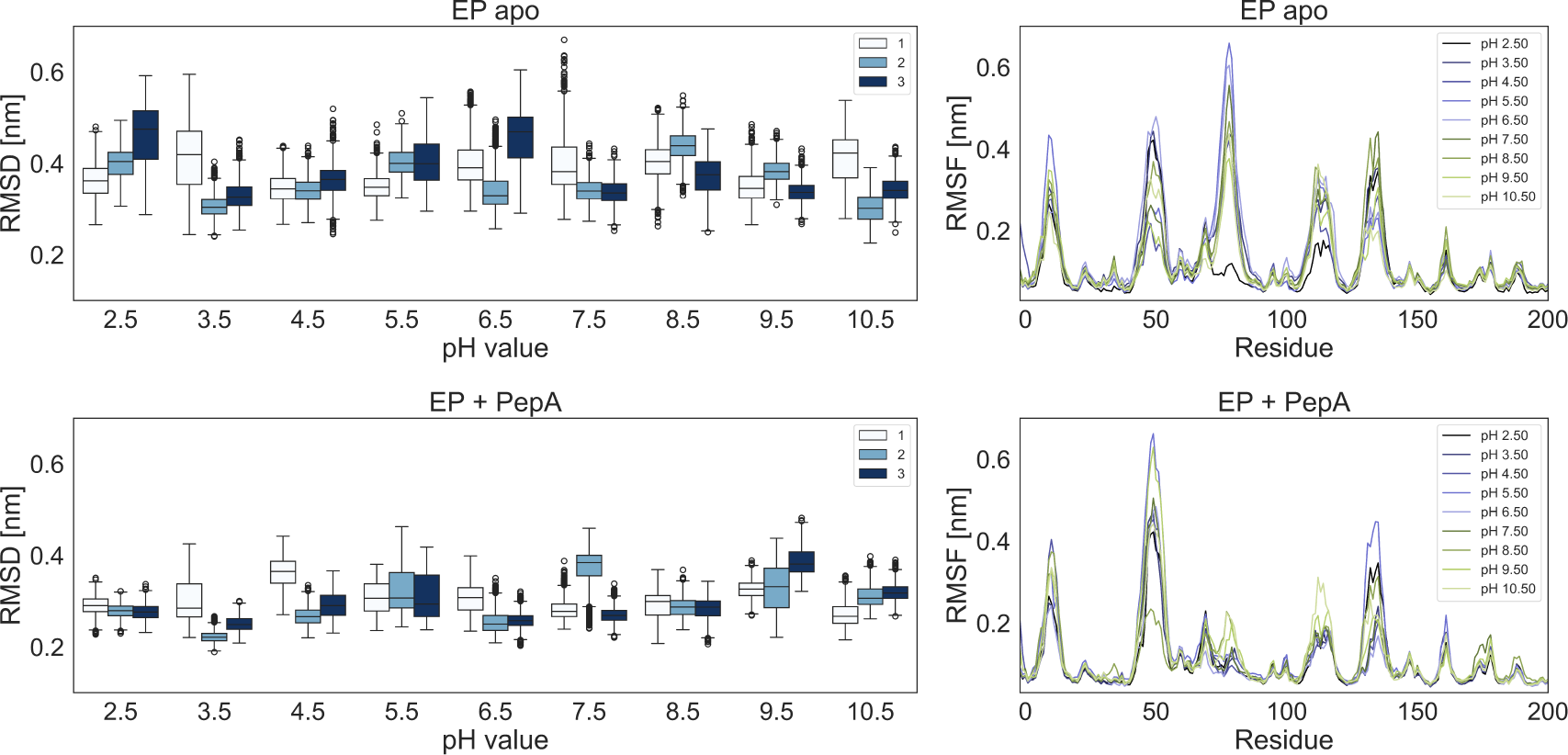
*Left* : Root mean square deviations (RMSD) of EP alone (”EP apo”) and in presence of pepA (”EP + PepA”) at different pH values, with respect to the crystal structure. Labels show the simulation repeat. *Right* : Averaged root mean square fluctuations (RMSF) of the C*_α_* of each residue at different pH values.

Focusing further on C*_α_* fluctuations (RMSF), we observed that the five top regions with the largest contribution to the RMSF correspond to loop-related motions, with no significant change on structured regions. In the pepstatin-bound structure, the motion of the loop covering residues 70-90 (commonly known as the binding flap, [19]) is strongly reduced, likely due to the stabilization with the binding partner. Moreover, we found that there are no large scale conformational motions that show dependence to the pH conditions.

In the previous section, our results suggested that a protonation event should take place upon binding, and that it should be connected to the aspartic acid residues. Thus, we evaluated the protonation propensities at different pH values for the aspartates within the binding site (i.e. Asp33, Asp35, Asp81 and Asp219), as they were the ones closer to pepA. With the protonation propensities, we obtained theoretical deprotonation curves and computed the theoretical p*K*_a_ (Figure 10).

**Figure 10:**
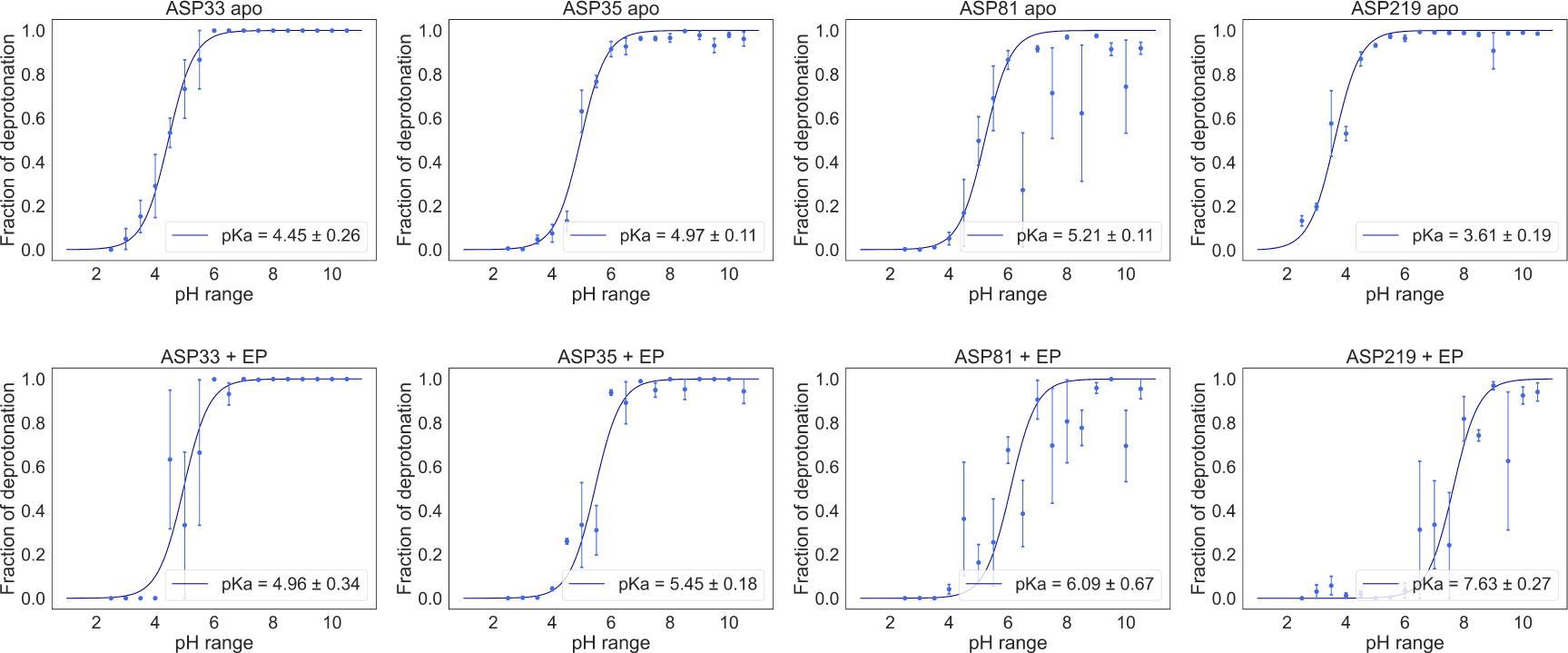
Titration curves of EP obtained from constant pH MD simulations for the single protein (top) and with pepA (bottom). For each of the four selected residues in this protein, the dots show the fraction of conformations in which the residue was deprotonated. Errors were estimated from the standard error of the mean for the three different replicas. The lines show the best fits to the Henderson–Hasselbalch equation. The p*K*_a_ values for each titratable residue are listed.

Indeed, the fraction of deprotonation curves suggest that the binding event shifts up the p*K*_a_ of all of the identified aspartates. Moreover, we can observe a high shift in the case of Asp219, from a p*K*_a_ of 3.6 to a p*K*_a_ of 7.6. This would also support the previous findings: that at pH 4.6 and 7.0, there is a change on total protonation (Δ*n*_H+_) close to one. At higher pH values such as pH 7.6, this effects become less prominent.

## 4 Conclusions

We were able to detect a proton transfer event for the complexation of pepA to EP. This was found in ITC experiments which revealed an overall proton uptake of 1.67 ± 0.12 moles of protons. This deviates from a published study [12] and can be explained by differences in the experimental setup. However, both ITC studies (our own and the published one [12]) were run at a pH of 7 which is significantly different from pH of 4.6 at which crystal structures exclusivey were determined so far.

This prompted us for follow-up studies such as a novel protein crystal structure determined at higher pH. Its analysis did not reveal any major conformational changes and therefore rules out its impact on the experimentally determined proton transfer event. Furthermore, this outcome justifies the usage of the already existing crystal structure (pH 4.6) as starting point for p*K*_a_ calcutions.

These calculations were performed in an implicit and explicit solvent manner and both agree in the structural interpretation of the proton transfer event. Additionally, constant pH molecular dynamics simulations show that the different pH conditions do not strongly affect the conformational dynamics of EP. This is in accordance with the novel crystal structures and underlines the power of this simulation technique. The findings from our study underline the necessity to crucially inspect possible proton transfer events. If those are not correctly accounted for the follow-up structure-based studies might easily evolve in an undesired direction.

## 5 Materials and Methods

### 5.1 Protein purification

The isolation of EP from Suparen (kindly provided by DSM Germany GmbH – Food Specialties, Düsseldorf) in 0.1 M NaOAc buffer pH 4.6 was performed as previously described by Köster *et al.*[20] For the experiments performed at a pH of 7.0, EP was dialyzed using the Slide-A-lyzer^®^ G2 cassette (ThermoFisher, Waltham, MA, USA) with a 10 kDa molecular weight cutoff. The dialysis was performed in a 600 mL beaker. The membrane of the cassette was soaked in the dialysis buffer for 2 min by holding it (with gloves) in the beaker filled with dialysis buffer at pH 7.0 (TRIS, ADA, HEPES and phosphate, respectively). After pre-soaking, the cassette was carefully dried with a paper towel to prevent buffer from entering the cassette, but without touching the membranes. 3 mL of at least 4 mg mL^−1^ protein solution was added to the cassette using a 200 µL micropipette. When the cassette was closed with the stopper, any remaining air was gently evacuated from the dialysis chamber. The protein was then dialyzed against 500 mL of dialysis buffer for 2 hours at RT with slow stirring. The buffer was then replaced with 500 mL of fresh buffer and stirred again for 2 hours at RT. After another change of buffer, the dialysate was stored in the refrigerator overnight (12-20 hours). The next day, the protein solution was removed using a syringe with a long cannula. The protein was stored on ice before being shock frozen in liquid nitrogen to determine the concentration using NanoDrop™2000c (Thermo Scientific, USA).

### 5.2 Protein crystallization

The protein was crystallized using an adapted protocol by Köster et al [20]. The protein was concentrated to 5 mg mL^−1^ and stored in 100 mM sodium acetate pH 4.6. Crystals of suitable size were grown in 4 µl drops consisting of 1:1 protein:reservoir solution, equilibrating against 1 mL of 100 mM sodium acetate pH 4.6, 100 mM ammonium acetate, 24 % PEG4000 on Cryschem plates (Hampton Research, USA) at 19°C. More reliable crystal growth could be achieved using streak seeding. Crystals suitable for data collection usually grew in 1-2 weeks. Co-crystallization with pepA was performed similarily, but solid pepA was first immobilized on the crystallization plates by evaporating 2 µL of a saturated pepA DMSO stock (¡100 mM) and then crystallization drops were set up on top of the solid. A soaking condition with pH 7.6 was prepared with a saturated concentration of pepA. Then, two crystals grown with pepA were soaked per condition for 24 hours. Crystals were then harvested from the drops with nylon loops and flash frozen with liquid nitrogen.

Data was collected at DESY P11[21]. The data was processed with XDS[22]. The structure was solved via molecular replacement and iterative cycles of model building in Coot[23] and refinement in Phenix[24] were performed.

### 5.3 Isothermal titration calorimetry

ITC measurements were performed using the MicroCal PEAQ-ITC Automated ITC (Malvern Panalytical, Worcestershire, UK). A 500 µM pepA solution containing 0.1 M of the different buffers (HEPES, TRIS, ADA, phosphate) of pH 7.0, 2.5 % (v/v) DMSO and 0.1 % (v/v) Tween20 was titrated into a 50 µM EP solution of the same buffer to a final stoichiometry of *N* = 2 (pepA:EP). For the reference titration, the same titrant was titrated into the buffer solution. The obtained thermogram peaks of all titrations were integrated and fitted with MicroCal PEAQ-ITC Analysis Software 1.41.

### 5.4 Implicit solvent p*K* _a_ calculations

The protein p*K*_a_ prediction was performed using a program based on the Zap finite difference PB solver by OpenEye, Cadence Molecular Sciences.[25, 1] As input, the workflow required a PDB file of the complex of interest and starting values for the pH and the p*K*_a_ of the ligand. The initial p*K*_a_ of pepA was predicted using ChemAxon’s Marvin p*K*_a_ plugin.[26] The ligand was then extracted from the original PDB file and hydrogens were added. Then, partial charges were assigned to both microspecies using the AM1BCC algorithm using the SZYBKI program by OpenEye, Cadence Molecular Sciences.[27] For the partial charges of the protein, Delphi radii and CHARMM36 all-hydrogen partial charges were used. An internal dielectric of 15, an ionic strength of 0.05 M and a pH of 7.0 were used. Reference p*K*_a_ values were as follows: Asp 4.0, His 6.5, Glu 4.5, Tyr 9.8, Lys 10.4 and Arg 12.5. Cys residues were not titrated and modeled in a neutral, protonated state. After protein and ligand were merged into one file and missing hydrogens were added to the protein, the processed microspecies, meaning the protonated and deprotonated form, were saved in a text file, which was used as input for subsequent steps. For the titratable amino acid residues listed above and the ligand, titration reactions were performed and p*K*_a_ values were calculated. All hydrogen atoms were explicitly modelled, and except for the orientation of the OH protons, which were sampled in 10° steps, the rest of the structure was static. Optimization of the ionization state and OH orientation was achieved by applying ten million Monte Carlo steps.[25]

For the p*K*_a_ calculation of the two aspartates in the catalytic dyad in the HIV-1 PR, the initial results were post-processed due to the electrostatically strongly coupled system: the two residues are in close proximity to each other at about 3Å and therefore behave more like a diprotic acid (e.g. maleic acid). This has already been described by Czodrowski *et al.*[28]

### 5.5 Constant pH molecular dynamics simulation

The initial structure of bound EP was taken from the PDB data bank (PDB ID: 4ER2 [29]) For the apo structure, the ligand was removed from the binding site manually. Excepting the aspartates, standard protonation states for all side chains were assumed through an in-house script. Both apo and pepstatin-bound structures were solvated in truncated octahedron water boxes with 1 nm between the protein and the boundaries of the box.

Constant pH molecular dynamics (CpHMD) simulations were performed using the GROMACS 2021-dev-beta implementation, where the constant pH approach was implemented by leveraging continuous lambda dynamics.[30] The proteins were represented using the implementation of Buslaev et. al. of CHARMM36m all-atom, due to the improved dihedral sampling in constant-pH simulations.[31] Missing parameters were obtained through the CHARMM Force Field server.[32] Ionic strength was set to 0.15 mM. The systems went through energy minimization, 1 ns heating with the V-rescale thermostat[33] to 300 K, 1 ns pressure coupling with the C-rescale barostat[34] to 1 atm. For the van der Waals interactions, a switching function of 10 to 12 Å was used. A real-space cutoff of 12 Å was applied in the electrostatics calculations with the particle-mesh Ewald method.

In the production run, the lambda dynamics is allowed, effectively exploring the protonation/deprotonation space of the selected residues at a given pH. In the CpHMD simulations, the side chains of the selected aspartates were allowed to titrate. The simulation of both apo EP and EP+pepstatin used 17 pH replicas in the pH range 2.5 to 10.5, with a spacing of 0.5 pH units, and 300 ns per pH-replica. Each system was simulated by triplicate to reduce dependency from the starting seed. The initial 50 ns from each replica were discarded in the analysis.

## 6 Acknowledgements

We are grateful to Luca Kröll and Sascha Jung for their valuable contributions. Parts of the results of this study were acquired with an ITC device funded by the Major Research Instrumentation Program of the German Research Foundation under grant No. INST 212/429-1 FUGG.

## References

[1] Honig B, Nicholls A. Classical electrostatics in biology and chemistry. Science. 1995;268:1144–1149

[2] Warshel A. Calculations of Enzymatic Reactions: Calculations of pKa, Proton Transfer Reactions, and General Acid Catalysis Reactions in Enzymes. Biochemistry. 1981;20:3167–3177

[3] Stanton C, Houk KN. Benchmarking pK prediction methods for residues in proteins. J Chem Theory Comput. 2008;4:951–966

[4] Neeb M, Czodrowski P, Heine A, Barandun LJ, Hohn C, Diederich F et al. Chasing protons: How isothermal titration calorimetry, mutagenesis, and pKa calculations trace the locus of charge in ligand binding to a tRNA-binding enzyme. J Med Chem. 2014;57(13), 5554–5565

[5] Hofer F, Kraml J, Kahler U, Kamenik AS, Liedl KR. Catalytic Site pKa Values of Aspartic, Cysteine, and Serine Proteases: Constant pH MD Simulations. J Chem Inf Model. 2020;60:3030–3042.

[6] Di Russo NV, Estrin DA, Martí MA, Roitberg AE. pH-Dependent Conformational Changes in Proteins and Their Effect on Experimental pKas: The Case of Nitrophorin 4. PLoS Comput Biol. 2012;8:88–100.

[7] Wlodawer A, Li M, Gustchina A, Dauter Z, Uchida K, Oyama H, et al. Inhibitor complexes of the Pseudomonas serine-carboxyl proteinase. Biochemistry 2001;40:15602–15611.

[8] Dostál J, Pecina A, Hrušková-Heidingsfeldová O, Marečková L, Pichová I, Řezáčová P, et al. Atomic resolution crystal structure of Sapp2p, a secreted aspartic protease from *Candida parapsilosis*. Acta Crystallogr Sect D Biol Crystallogr 2015;71:2494–2504.

[9] Schiebel J, Gaspari R, Sandner A, Ngo K, Gerber HD, Cavalli A et al. Charges Shift Protonation: Neutron Diffraction Reveals that Aniline and 2-Aminopyridine Become Protonated Upon Binding to Trypsin. Angew Chemie - Int Ed. 2017;56:4887–4890.

[10] Bastos M, Abian O, Johnson CM, Ferreira-da-Silva F, Vega S, Jimenez-Alesanco A, et al. Isothermal titration calorimetry. Nat Rev Methods Prim. 2023;3.

[11] Krimmer SG, Klebe G. Thermodynamics of protein-ligand interactions as a reference for computational analysis: How to assess accuracy, reliability and relevance of experimental data. J Comput Aided Mol Des. 2015;29:867–883.

[12] Gómez J, Freire E. Thermodynamic Mapping of the Inhibitor Site of the Aspartic Protease Endothiapepsin. J Mol Biol. 2015;252:337–350.

[13] Czodrowski P, Sotriffer CA, Klebe G. Protonation Changes upon Ligand Binding to Trypsin and Thrombin: Structural Interpretation Based on pKa Calculations and ITC Experiments. J Mol Biol. 2007;367:1347–1356.

[14] Goldberg RN, Kishore N, Lennen RM. Thermodynamic quantities for the ionization reactions of buffers. J Phys Chem Ref Data. 2002;31:231–370.

[15] Cubrilovic D, Zenobi R. Influence of dimehylsulfoxide on protein-ligand binding affinities. Anal Chem. 2013;85:2724–2730.

[16] Schiebel J, Radeva N, Krimmer SG, Wang X, Stieler M, Ehrmann FR, et al. Six Biophysical Screening Methods Miss a Large Proportion of Crystallographically Discovered Fragment Hits: A Case Study. ACS Chem Biol 2016;11:1693–1701.

[17] Hyland LJ, Tomaszek TA, Meek TD. Human Immunodeficiency Virus-1 Protease. 2. Use of pH Rate Studies and Solvent Kinetic Isotope Effects To Elucidate Details of Chemical Mechanism. Biochemistry. 1991;30:8454–8463.

[18] Henderson JA, Harris RC, Tsai CC, Shen J. How Ligand Protonation State Controls Water in Protein-Ligand Binding. J Phys Chem Lett. 2018;9:5440–5444.

[19] Hong L, Tang J. Flap Position of Free Memapsin 2 (β-Secretase), a Model for Flap Opening in Aspartic Protease Catalysis. Biochemistry. 2004;43:4689–4695.

[20] Köster H, Craan T, Brass S, Herhaus C, Zentgraf M, Neumann L, et al. A small nonrule of 3 compatible fragment library provides high hit rate of endothiapepsin crystal structures with various fragment chemotypes. J Med Chem. 2011;54:7784–7796.

[21] Meents A, Reime B, Stuebe N, Fischer P, Warmer M, Goeries D, et al. Development of an in-vacuum x-ray microscope with cryogenic sample cooling for beamline P11 at PETRA III. X-Ray Nanoimaging: Instruments and Methods. 2013 8851:88510K.

[22] Kabsch W (2010) XDS. Acta Crystallogr Sect D. 2010;66:125–132.

[23] Emsley P, Lohkamp B, Scott WG, Cowtan K. Features and development of Coot. Acta Crystallogr Sect D Biol Crystallogr. 2010;66:486–501.

[24] Liebschner D, Afonine PV, Baker ML, Bunkoczi G, Chen VB, Croll TI, et al. Macromolecular structure determination using X-rays, neutrons and electrons: Recent developments in Phenix. Acta Crystallogr Sect D Struct Biol 2019;75:861–877.

[25] Word JM, Nicholls A. Application of the Gaussian dielectric boundary in Zap to the prediction of protein pKa values. Proteins Struct Funct Bioinforma 2011;79:3400–3409.

[26] MarvinSketch 19.10, ChemAxon, www.chemaxon.com, 2019.

[27] SZYBKI 2.7.1.0. OpenEye, Cadence Molecular Sciences, Santa Fe, NM, 2024.

[28] Czodrowski P, Sotriffer CA, Klebe G. Atypical protonation states in the active site of HIV-1 protease: A computational study. J Chem Inf Model. 2007;47:1590–1598.

[29] Pearl L, Blundell T. The active site of aspartic proteinases. FEBS Lett. 1984;174:96–101.

[30] Aho N, Buslaev P, Jansen A, Bauer P, Groenhof G, Hess B. Scalable Constant pH Molecular Dynamics in GROMACS. J Chem Theory Comput. 2022;18:6148–6160.

[31] Buslaev P, Aho N, Jansen A, Bauer P, Hess B, Groenhof G. Best Practices in Constant pH MD Simulations: Accuracy and Sampling. J Chem Theory Comput. 2022;18:6134–6147.

[32] Vanommeslaeghe K, Hatcher E, Acharya C, Kundu S, Zhong S, Shim J, et al. CHARMM general force field: A force field for drug-like molecules compatible with the CHARMM all-atom additive biological force fields. J Comput Chem. 2010;31:671–690.

[33] Bussi G, Donadio D, Parrinello M. Canonical sampling through velocity rescaling. J Chem Phys. 2007;126.

[34] Bernetti M, Bussi G. Pressure control using stochastic cell rescaling. J Chem Phys. 2020;153:1–11.

